# Identification of hidden population structure in time-scaled phylogenies

**DOI:** 10.1101/704528

**Authors:** Erik M. Volz, Carsten Wiuf, Yonatan H. Grad, Simon D.W. Frost, Ann M. Dennis, Xavier Didelot

## Abstract

Population structure influences genealogical patterns, however data pertaining to how populations are structured are often unavailable or not directly observable. Inference of population structure is highly important in molecular epidemiology where pathogen phylogenetics is increasingly used to infer transmission patterns and detect outbreaks. Discrepancies between observed and idealised genealogies, such as those generated by the coalescent process, can be quantified, and where significant differences occur, may reveal the action of natural selection, host population structure, or other demographic and epidemiological heterogeneities. We have developed a fast non-parametric statistical test for detection of cryptic population structure in time-scaled phylogenetic trees. The test is based on contrasting estimated phylogenies with the theoretically expected phylodynamic ordering of common ancestors in two clades within a coalescent framework. These statistical tests have also motivated the development of algorithms which can be used to quickly screen a phylogenetic tree for clades which are likely to share a distinct demographic or epidemiological history. Epidemiological applications include identification of outbreaks in vulnerable host populations or rapid expansion of genotypes with a fitness advantage. To demonstrate the utility of these methods for outbreak detection, we applied the new methods to large phylogenies reconstructed from thousands of HIV-1 partial *pol* sequences. This revealed the presence of clades which had grown rapidly in the recent past, and was significantly concentrated in young men, suggesting recent and rapid transmission in that group. Furthermore, to demonstrate the utility of these methods for the study of antimicrobial resistance, we applied the new methods to a large phylogeny reconstructed from whole genome *Neisseria gonorrhoeae* sequences. We find that population structure detected using these methods closely overlaps with the appearance and expansion of mutations conferring antimicrobial resistance.

---

Quantifying the role of population structure in shaping genetic diversity is a longstanding problem in population genetics. When information about how lineages are sampled is available, primarily geographic location, a variety of statistics are available for describing the magnitude and role of population structure (Hartl et al. 1997). In pathogen phylogenetics, such geographic ‘meta-data’ has been instrumental in enabling the inference of transmission rates over space (Dudas et al. 2017), host species (Lam et al. 2015), and even individual hosts (De Maio et al. 2018). Population structure shapes genetic diversity, but can the existence of structure be inferred directly from genetic data in the absence of structural covariates associated with each lineage, such as if the geographic location or host species of a lineage is unknown?

The problem of detecting and quantifying such ‘cryptic’ population structure has become a pressing issue in several areas of microbial phylogenetics. For example, in bacterial population genomics studies, a wide diversity of methods have been recently developed to classify taxonomic units based on distributions of genetic relatedness (Mostowy et al. 2017; Tonkin-Hill et al. 2019, 2018; Beugin et al. 2018). In a different domain, pathogen sequence data have been used for epidemiological surveillance, and ‘clustering’ patterns of closely related sequences have been used to aid outbreak investigations and prioritise public health interventions (Eyre et al. 2012; Dennis et al. 2014; Miller et al. 2014; Ledda et al. 2017). In both population genomics studies and outbreak investigations, a common thread is the absence of variables about sampled lineages that can be correlated with phylogenetic patterns. For example, in outbreak investigations, host risk behaviour and transmission patterns are not usually observed and must be inferred. It is not known a priori which clades are more or less likely to expand in the future, although there is active research addressing this problem, such as to predict the emergence of strains of influenza A virus (Klingen et al. 2018) or to forecast the effect of antibiotic usage policies on the prevalence of resistant variants (Whittles et al. 2017).

In time-scaled phylogenies, the effects of population structure often appear as a difference in the distribution of branch lengths in clades circulating in different populations (Dearlove and Frost 2015). Figure 1 shows a simulated genealogy from a structured coalescent process (Notohara 1990). In two clades, the effective population size grows exponentially, and in the remaining clade, the effective size remains constant. Consequently, the number of lineages through time show noticeably different patterns of relatedness. For the clades with growing size, most coalescent events occur in the distant past when the size was small.

**Figure 1:**
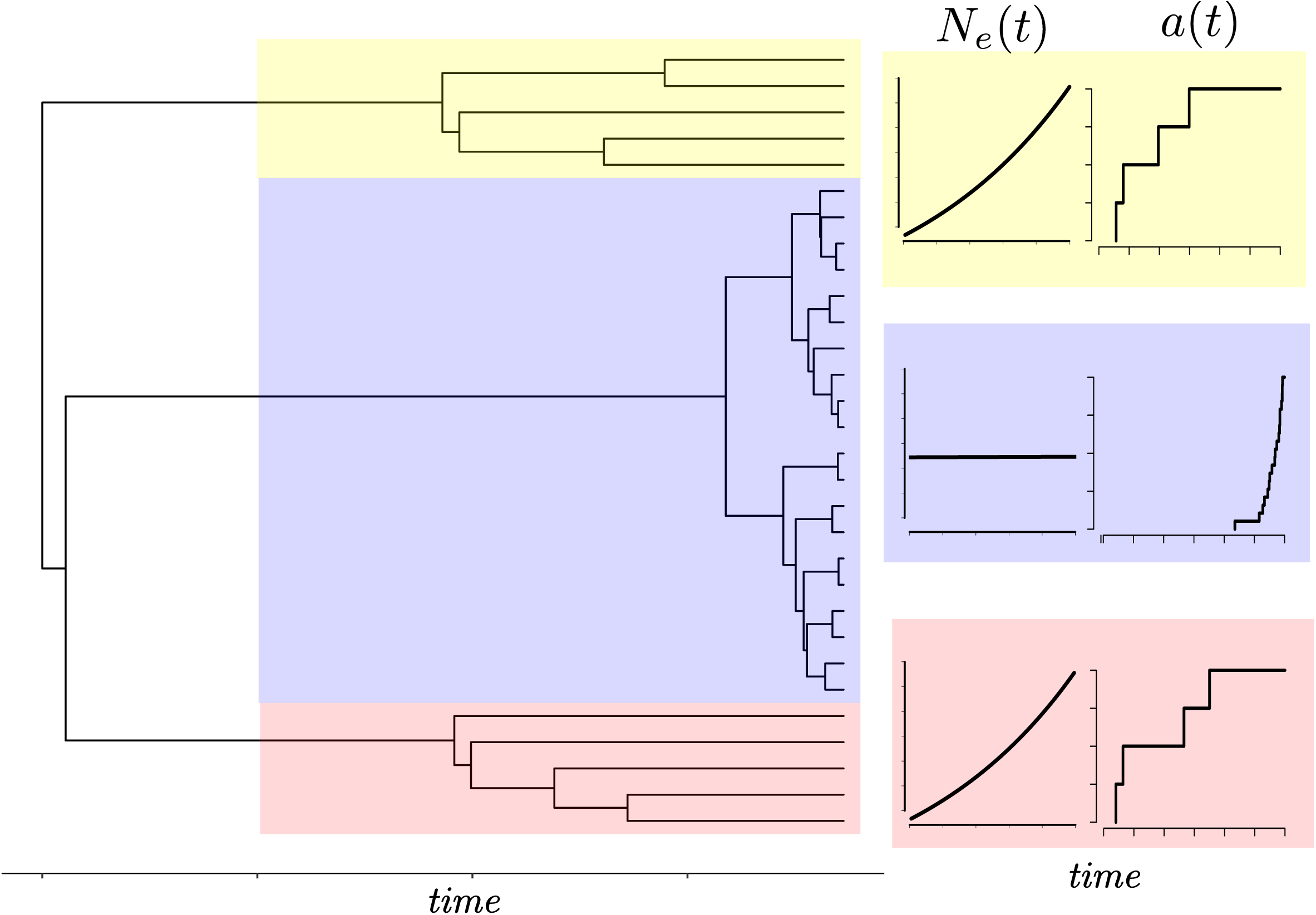
A genealogy simulated from a structured coalescent process with two demes, one of which has constant effective population size (clade highlighted in blue), and the other having effective population size growing exponentially (clades highlighted in red and yellow). Migration of lineages occurs at a small constant rate in one direction from the constant size deme to the growing deme. The corresponding plots at the right show a caricature of the effective population size and number of lineages through time in each clade.

Supposing that the deme from which lineages were sampled was not observed, it is clear from visual inspection of Figure 1 which lineages were sampled from a growing population. Nevertheless, there is a paucity of objective methods readily available to automate the process of identifying temporally distinct clades. This process cannot be done manually when the differences in distributions are less obvious, and needs to be based on a theoretically grounded statistical test. Furthermore, in Figure 1, the red and yellow clades are distantly related. Their most recent common ancestor (MRCA) is at the root of the tree, but they have a very similar distribution of coalescent times suggesting that they were generated by similar demographic or epidemiological processes. For example, this can happen in infectious disease epidemics, when lineages independently colonise the same host population with greater susceptibility or higher risk behaviour (Dearlove et al. 2017). It is therefore also desirable to have an automated method for identifying polyphyletic taxonomic groups defined by shared inferred population histories as opposed to genetic or phenotypic traits.

Here we develop a statistical test for detecting if clades within a time-scaled genealogy have evidence for unobserved population structure. Our approach is to develop a statistic based on an unstructured coalescent process. This allows us to test a null hypothesis that two clades are both generated by the same coalescent process. In this case, the coalescent model provides a theoretical prediction of the order of the coalescent times between the two clades in the absence of population structure. On the basis of this statistical test, we also develop algorithms for systematically exploring possible partitions of a genealogy into distinct sets representing evolution within latent populations with different demographic or epidemic histories. Notably, these algorithms not only allow us to detect outlying clades with very different genealogical patterns, but also to find and classify distantly related clades which likely have similar demographic or epidemic histories.

## Materials and Methods

As a starting point for our methodology, we assume a time-scaled phylogeny has been estimated from genetic data, for example using one of the recently developed fast methods (To et al. 2016; Volz and Frost 2017; Didelot et al. 2018; Sagulenko et al. 2018; Tamura et al. 2018; Miura et al. 2019). Alternatively, summary trees obtained from full Bayesian approaches as implemented in BEAST (Suchard et al. 2018; Bouckaert et al. 2014) or RevBayes (Höhna et al. 2016) can be used, although these typically incorporate population genetic models which presume a particular form of population structure or a lack of population structure. Some precise terminology and notation is required related to the structure of these time-scaled trees since the basis of our approach concerns comparisons between different subsets of the tree.

### Notation

The tree has *n* terminal nodes (nodes with no descendants), is rooted, and is bifurcating (there are *n* − 1 internal nodes each with exactly two descendants). Being rooted implies there is one node with no ancestor. Mathematically we describe this tree as a node-labelled directed acyclic graph:

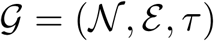

where 𝒩 is a set of 2*n* − 1 nodes, *ε* ⊆ {(*u, v*)|*u, v* ∈ 𝒩^2^} is the set of 2*n* − 2 edges or ‘lineages’, and *τ*: 𝒩 → ℝ_≥0_ defines the time of each node. With reference to an edge (*u, v*) ∈ *ε* we say that *u* is the ‘direct ancestor’ and *v* is the ‘direct descendant’ and we require *τ* (*u*) *< τ* (*v*). Nodes are further classified into two sets: ‘tips’ (terminal nodes) denoted 𝒯 with no descendants and internal nodes denoted ℐ with exactly two direct descendants. The trees may be heterochronous, meaning that tips of the tree can represent samples taken at different time points.

For a node *u* ∈ 𝒩 we define the clade *C*_*u*_ to be the set of nodes descending from *u*, that is, the node *u* and all *v* ∈ 𝒩 such that there is a directed path of edges from *u* to *v*. We say that nodes *v* in *C*_*u*_ are ‘descended from’ *u*. We will also have occasion to define clades ‘top down’ in terms of a subset of tips in the tree. For this, we define the most recent common ancestor MRCA(*X*) of a set *X* ⊆ 𝒯 to be the most recent node *u* such that *X* ⊆ *C*_*u*_, that is, all other nodes *v* with *X* ⊆ *C*_*v*_ have *τ* (*v*) *< τ* (*u*). Then we let the top-down clade *B*_*X*_ be defined as

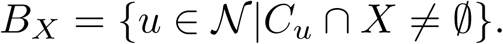

Note that *B*_*X*_ includes the tips *X* as well as some nodes ancestral to MRCA(*X*).

In general *B*_*X*_ ≠ *C*_MRCA(*X*)_ since *X* does not necessarily include all tips descending from MRCA(*X*). We will also need to refer to the nodes corresponding to coalescent events among lineages of the set *X* only, excluding those between lineages of *X* and lineages of the complement of *X*,

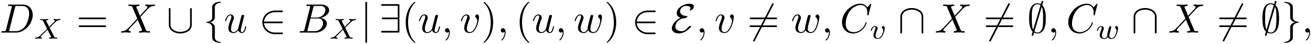

Figure 2A illustrates a tree and the sets *B*_*X*_, *D*_*X*_, and *C*_MRCA(*X*)_.

**Figure 2:**
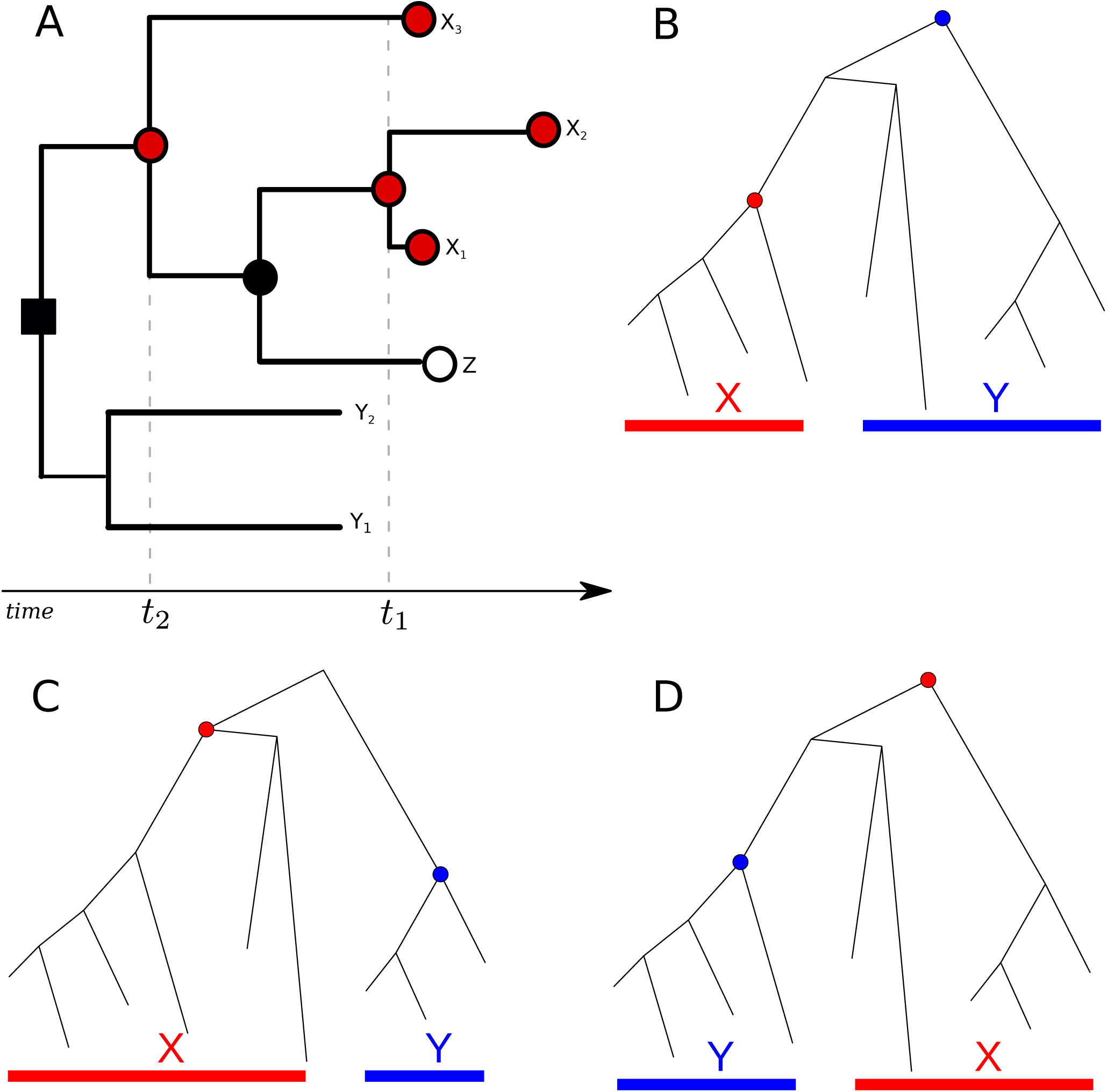
Coalescent trees for illustrating taxonomic relationships and notation used throughout the text. In panel A, the shape and colour of nodes correspond to variables *B*_*X*_, *D*_*X*_, and *C*_MRCA(*X*)_ in relation to the set of tips *X* = {*x*_1_, *x*_2_, *x*_3_}. All circles regardless of colour correspond to *C*_MRCA(*X*)_. All filled shapes (red or black, square or circle) correspond to *B*_*X*_. Note that this includes nodes ancestral to the MRCA of *X*. All red filled circles correspond to *D*_*X*_. Two coalescent events occur among nodes in *D*_*X*_ at times *t*_1_ and *t*_2_. Panels B-D show a coalescent tree and examples of potential taxonomic relationships between two clades. Prior knowledge of taxonomic relationships between *X* and *Y* influences the probability that the next coalescent event will be observed in clade *X*.

Since each node has a time, we can define the set of ‘extant’ lineages 𝒜(*t*) at a particular time *t* to be the set of nodes occurring after time *t* with a direct ancestor before time *t*,

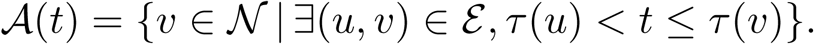

We might also refer to the number of extant lineages at time *t, a*(*t*) = |𝒜(*t*)|, and if considering the number of extant lineages within a particular clade ancestral to (and including) *X* we write

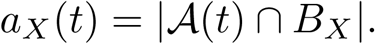

### Non-parametric test for a given pair of clades

With the above notation, the rank-sum statistic can now be defined which will form the basis for subsequent statistical tests and can be used to compare any pair of clades in the tree.

Let *X* and *Y* represent disjoint sets of tips as represented in Figure 2B-D. Having sorted the nodes according to time and assigned a corresponding rank to each internal node, this statistic computes the sum of ranks in a given clade in comparison to a different clade:

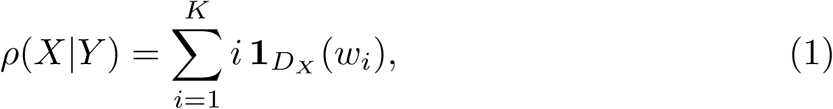

where *w*_*i*_ is an element of *S*_*X,Y*_ = (*w*_1_, *w*_2_, …, *w*_*K*_) which is the sequence of internal nodes in *D*_*X*_ ∪ *D*_*Y*_ sorted by time (present to past). And, **1**_*A*_(*u*) is an indicator that takes the value 1 if *u* ∈ *A* and is zero otherwise. Note that *ρ*(*X*|*Y*) is asymmetric in *X* and *Y*. Also note that *ρ*(*X*|*Y*) makes use of *D*_*X*_ and *D*_*Y*_, not *B*_*X*_ and *B*_*Y*_, because we are interested in the relative ordering of coalescent events among lineages of *X* and *Y*. Although the statistic is defined for all sets disjoint sets *X* and *Y* the examples we consider below apply to the case that the intersection of *D*_*X*_ and *D*_*Y*_ is empty. Only the ordering of the events matter, the absolute times are immaterial to the test.

Under a neutral coalescent process, the distribution of coalescent times in two clades ancestral to *X* and *Y* will depend on the number of extant lineages through time in both clades and on the effective population size *N*_*e*_(*t*) (Wakeley 2009). However, the distribution of the relative ordering of coalescent times only depends on the sizes of the clades. This distribution can be computed rapidly by Monte-Carlo simulation as shown below, provided that we know the probability that the next coalescent will be in *X* or *Y* as a function of the number of lineages ancestral to *X* and *Y*, given by *a*_*X*_ (*t*) and *a*_*Y*_ (*t*). We here provide new theoretical results on the distribution of the relative ordering of coalescence times under the null hypothesis that both *B*_*X*_ and *B*_*Y*_ are clades within a single tree generated by a neutral unstructured coalescent process. In the following we consider three different scenarios.

#### Event *E*_1_

Suppose that a clade *B*_*X*_ has a MRCA before any tip of *X* shares a common ancestor with the clade of another set of tips *Y*, disjoint to *X*. After lineages in *X* have found a common ancestor, the MRCA of *X* may or may not coalesce with lineages in *B*_*Y*_ before *Y* has found a common ancestor. Figure 2B-C illustrates trees that satisfy this condition. Note that in Figure 2B, a lineage in *Y* coalesces with the MRCA of *X* before lineages in *Y* find a MRCA and in Figure C, both *X* and *Y* have a common ancestor before they find a common ancestor with one another.

Observing a taxonomic pattern such as shown in Figure 2B-C is a random event in a stochastic unstructured coalescent process, and we denote this event by *E*_1_ (suppressing *X* and *Y* for convenience). Wiuf and Donnelly (Wiuf and Donnelly 1999) showed that the probability of observing *E*_1_, given the state of the tree at a particular time *t*, only depends on the number of lineages *z* = *a*_*X*_ (*t*) and *w* = *a*_*Y*_ (*t*),

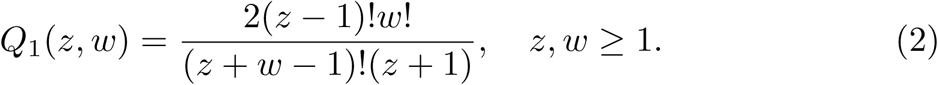

The numbers of extant lineages in *B*_*X*_ (or its complement) following each coalescent event conditional on *E*_1_ is a Markov chain. The transition probabilities of this chain are exactly those needed to simulate the null distribution of the test statistic *ρ*(*X*|*Y*). The probability that the next coalescent event is among lineages in the clade *B*_*X*_ given *E*_1_ (starting at a particular time *t*) was found by Wiuf and Donnelly (Wiuf and Donnelly 1999):

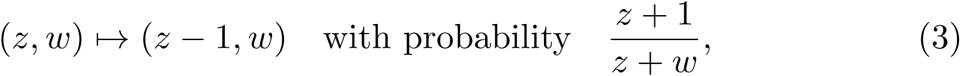

where the ancestral number of lineages of *X* and *Y* at time *t* are respectively *z* and *w*.

#### Event *E*_2_

We further derive analogous probabilities under slightly different conditions. Suppose we have disjoint sets of tips, *X* and *Y*. Let all lineages in *X* share a common ancestor before any share a common ancestor with *Y and* vice versa, all lineages in *Y* share a common ancestor before any share a common ancestor with tips in *X*. Figure 2C illustrates a tree and two clades that satisfy this condition, which we denote by *E*_2_. As before, the number of ancestors in *B*_*X*_ and *B*_*Y*_ will form a Markov chain, conditional on *E*_2_.

The probability that the next coalescent event is among lineages in the clade *B*_*X*_ given *E*_2_ at a particular time *t* and the current ancestral number of lineages of *X, z* = *a*_*X*_ (*t*), and *Y, w* = *a*_*Y*_ (*t*), can be given as:

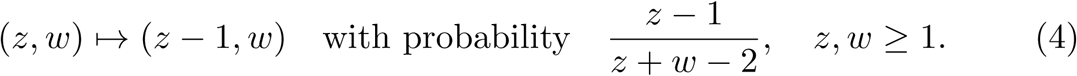

To see this, note that without conditioning on *E*_2_, the probability that the next coalescent is among ancestral nodes in *B*_*X*_ is

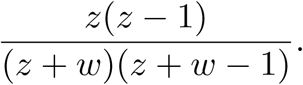

This is simply the ratio of the coalescent rate in *B*_*X*_, which is 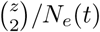, to the rate in *B*_*X*_ ∪ *B*_*Y*_, which is 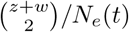. The effective population size is homogenous through the tree by hypothesis of the statistical test, and it cancels out in this ratio. The probability that the coalescent event would be between the clades ancestral to *X* and *Y* would be

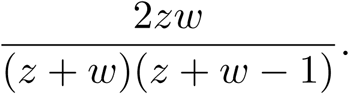

Event *E*_2_ has probability *Q*_2_(*z, w*), which must fulfil the recursion

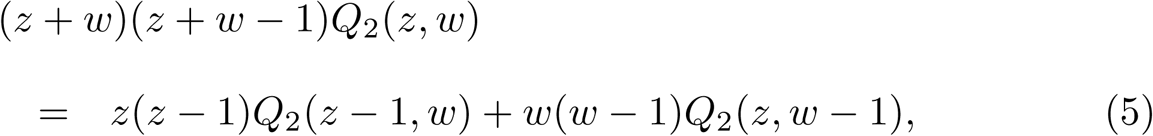

where *z, w* ≥ 1. If there is exactly one lineage in both *B*_*X*_ and *B*_*Y*_, then *Q*_2_(1, 1) = 1. If there is one lineage remaining in *B*_*X*_ and *w >* 1 in *B*_*Y*_, then *Q*_2_(1, *w*) is the probability that the next *w* − 1 coalescent events only occur between lineages in *B*_*Y*_ and do not include the single lineage ancestral to *X*. The probability of the next coalescent event being in *B*_*Y*_ is the probability of not selecting the *B*_*X*_ lineage when sampling two extant lineages without replacement:

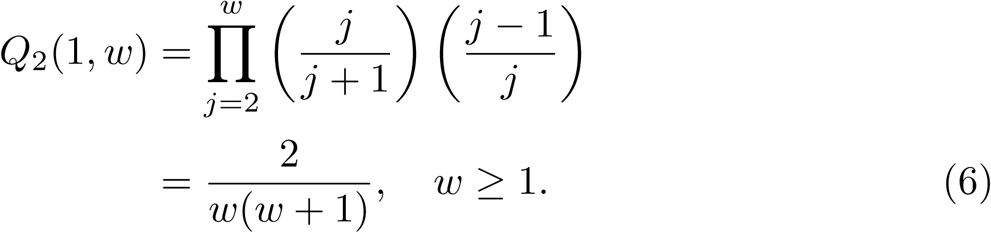

Similarly, 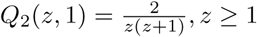. This recursion can be solved explicitly to give

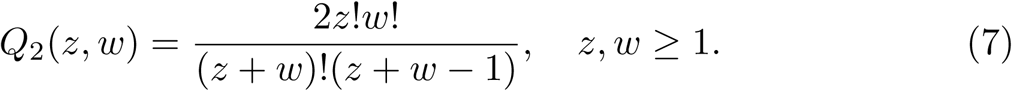

Now the transition probability (Equation 4) can be defined in terms of the rate of coalescence in *B*_*X*_ and *B*_*Y*_ and the probability of *E*_2_ being satisfied following the coalescent event:

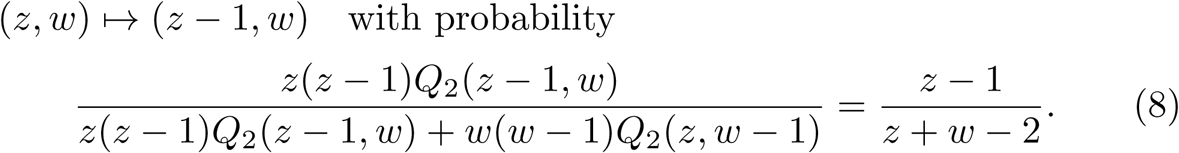

#### Event *E*_3_

Finally, we consider an event that is the union of events *E*_1_ and *E*_2_. We denote *E*_3_ to be the event that all *X* have a MRCA before sharing a common ancestor with lineages of *Y* and/or all lineages in *Y* have a MRCA before sharing an ancestor with lineages of *X*. All trees in Figure 2B-D satisfy this condition.

The probability of the event *E*_3_ can be defined in terms of *Q*_1_ and *Q*_2_ given previously:

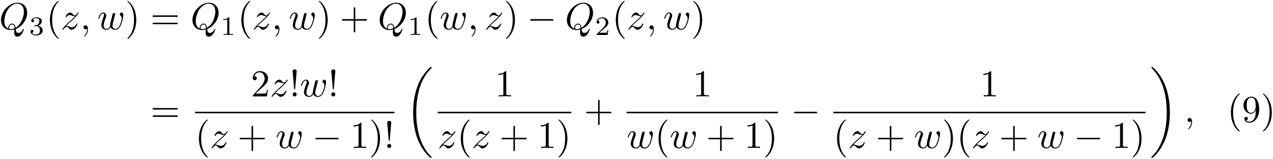

with *z* = *a*_*X*_ (*t*) and *w* = *a*_*Y*_ (*t*) being sample sizes at a particular time *t*, as before. The function *Q*_3_ satisfies the same recursion as above (Equation 5) with slightly different boundary conditions:

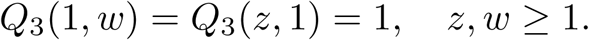

Transition probabilities can be derived as above by substituting *Q*_3_ for *Q*_2_ in Equation 8. The probability that the next coalescent event is among lineages in *D*_*X*_ conditional on *E*_3_ is

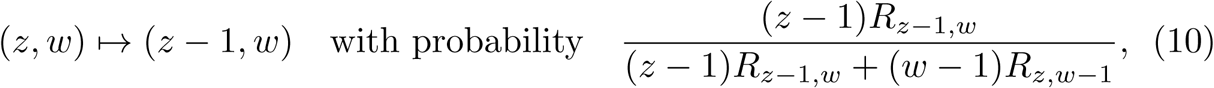

where

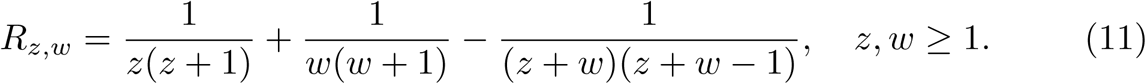

### Algorithms for detecting population structure

The null distribution of the test statistic *ρ*(*X, Y*) can be computed by Monte-Carlo simulation using Equations 3, 4 or 10 depending on the taxonomic constraints to be conditioned on. This can be computed given any pair of disjoint clades *X* and *Y*. Algorithm 1 in the Supplementary Material provides the simulation procedure for computing the two-sided p-values of an empirical measurement 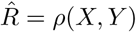, and we denote these p-values *ξ*(*X, Y, R*). The algorithm works by simulating many replicates of the rank-sum statistic conditional on the sets *X, Y*, and the taxonomic relationship between these clades. Furthermore, the order of sampling events and coalescent events is part of the data within a time-scaled phylogeny. Thus the simulation procedure does not simulate coalescent trees per se, but rather the number of lineages through time *a*_*X*_ (*t*) and *a*_*Y*_ (*t*) by proceeding from the most recent sample back to the MRCA of clades *X* and *Y*. Upon visiting a node in the ordered sequence of coalescent events, the algorithm selects at random a clade *D*_*X*_ or *D*_*Y*_ for this event using the transition probabilities from Equations 3, 4 or 10. Upon visiting a coalescent event, *a*_*X*_ (*t*) or *a*_*Y*_ (*t*) is incremented using the observed clade membership of the sample at that time. The end result of this simulation procedure is a large set of replicate rank-sum statistics which serves as a null distribution for comparison with the value computed from the time-scaled phylogeny.

While in principle this test allows comparison of any pair of disjoint clades, the number of possible comparisons is vast, and deriving a useful summary of taxonomic structure requires additional heuristic algorithms. These algorithms are designed to stratify clades into self-similar sets and to do so in a computationally efficient manner. Algorithm 2 in the Supplementary Material identifies ‘cladistic outliers’, which are clades that have a coalescent pattern that is different from the remainder of the tree. It performs a single pre-order traversal of the tree and greedily adds clades to the partition with the most outlying values of the test statistic. At each node *u* visited in pre-order traversal, Algorithm 2 examines all descendants *v* in *C*_*u*_ and compares *C*_*v*_ with to *C*_*u*_ \ *C*_*v*_. If no outliers are found, the algorithm will desist from searching *C*_*u*_ and the set of tips *C*_*u*_ ∩ *𝒯* will be added to the partition. If at least one outlier is found in *C*_*u*_, a search will begin on the biggest outlier (smallest p-value computed using Algorithm 1). The final result of this algorithm is a partition of *m* non-overlapping clades *M* = {*X*_1_, …, *X*_*m*_}.

In practice, it is often desirable to not compare very small clades against one another or much larger clades, so additional parameters are available to desist the pre-order traversal upon reaching a clade with few descendants. It is also often of practical interest to only compare clades that overlap in time to a significant extent, so yet another parameter is available to desist from comparing a pair of clades if few lineages in the pair ever coexist at any time.

Additional algorithms are required to detect polyphyletic relationships as depicted in Figure 1 which arise if, for example, distantly related lineages colonise the same area and have similar population dynamics or if near-identical fitness-enhancing mutations occur independently on different lineages. Figure 1 depicts two distantly related clades (yellow and red) with similar population dynamics, and it is desirable to classify these as a single deme based on shared population dynamic history. Algorithm 2 will partition tips of the tree into distinct clades with monophyletic or paraphyletic relationships, however an approach based on pre-order traversal of the tree can not on its own arrive at a polyphyletic partition of the tree. Therefore we can implement a final hierarchical clustering step in order to group similar clades as follows:

1. For each distinct pair of clades *X* and *Y* in partition *M*, compute 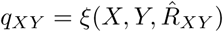.
2. Convert the p-value into a measure of distance between all clades: *d*_*XY*_ = |*F* ^−1^(*q*_*XY*_)|, where *F* ^−1^ is the inverse Gaussian cumulative distribution function (quantile function). Set *d*_*XX*_ = 0 for all *X*.
3. Perform a conventional hierarchical clustering using a threshold distance *F* ^−1^(1 − *α/*2) for confidence level *α*. Various clustering algorithms can be used at this point, and our software has implemented the ‘complete linkage’ algorithm (Everitt et al. 2001).

Algorithms 1 and 2 as well as the final hierarchical clustering step are implemented as an open source R package called *treestructure* available at https://github.com/emvolz-phylodynamics/treestructure. The R package supports parallelisation and includes facilities for tree visualisation using the *ggtree* package (Yu et al. 2017). The package provides convenience functions to output cluster and partition assignment for downstream statistical analysis in R.

### Simulation studies

To evaluate the potential for *treestructure* to detect outbreaks we applied the new method to phylogenies estimated from newly simulated data using a structured coalescent model as well as previously published simulation data based on a discrete-event branching process (McCloskey and Poon 2017). We also simulated trees and sequence data under a Kingman coalescent process to examine the distribution of the test statistic under the null hypothesis and to assess how statistical power of the test depends on sample size and the differences between clades.

The structured coalescent simulation was based on a model with two demes: a large deme with constant effective population size and a smaller deme which grows exponentially up to the time of sampling. Migration occurs at a constant rate in both directions between the growing and constant-size demes, and equal proportions of these two demes are sampled. Coalescent simulations were implemented using the *phydynR* package http://github.com/emvolz-phylodynamics/phydynR. All genealogies simulated from this model were comprised of 1000 tips with 200 of these sampled from the growing deme. Each of 100 simulations were based on different parameters such that there was a spectrum of difficulty identifying population structure from the trees. The sample proportion was chosen uniformly between 5% and 75% and, the growth rate in the growing deme was chosen uniformly between 5% and 100% per year. Bidirectional migration between demes was fixed at 5% per year. While most tips were sampled at a single time point, 50 tips from the constant-size deme were distributed uniformly through time in order to facilitate molecular clock dating. Multiple sequence alignments were simulated based on trees using seq-gen (Rambaut and Grass 1997). Each sequence comprised 1000 nucleotides from a HKY model with a substitution rate of 10^−3^ per site per year, which is a typical value for RNA viruses. A neighbor joining tree was estimated from each alignment and dated phylogenies estimated using the *treedater* R package (Volz and Frost 2017) with a strict molecular clock. The *treestructure* algorithm was applied to each phylogeny using the default *α* = 1% threshold.

In order to test the specificity of our method, we also simulated 1,000 trees under an unstructured Kingman coalescent process using the *rcoal* function in the *ape* R package version 5.2. These trees each had 50 tips and an effective population size of 0.025. Sequence data and neighbor joining trees were generated as described above. The *estimate*.*dates* command (Jones and Poon 2016) in the *ape* R package version 5.2 was used to estimate time-scaled trees. The *treestructure* algorithm was applied to both the coalescent trees and to the trees estimated based on the simulated sequences. The test statistic was tabulated for each clade size from 5 to 45 leading to approximately 10,000 observations of the test statistic in total, and about 250 observations for each clade size.

A further set of Kingman coalescent simulations was carried out to assess the statistical power of our method. We simulated paired coalescent trees of different sizes and with different effective population sizes, and each pair of coalescent trees was then joined at a common root. Branch lengths at the root node were adjusted to ensure the trees were ultrametric. One tree in each pair was small with 10, 20 or 40 tips, whereas the other had 200 tips. The *treestructure* algorithm was used to compute the normalized test statistic at the MRCA of the minority clade. The effective population size in the minority clade was varied to provide differing levels of contrast. Note that even if the effective population size is the same in the majority and minority clades, the topology of the combined tree may differ substantially from the Kingman model, so that the minority clade may be detected by the *treestructure* algorithm. To effectively ‘hide’ the structure caused by the construction of the combined trees, we can set the effective population size of the minority clade to be *zN*_*e*_*/w* where *z* is the number of tips in the minority tree, *w* is the number of tips in the majority tree, and *N*_*e*_ is the effective size of the majority tree. By doing so, the initial coalescent rate in both trees will be as expected under the Kingman model for the combined tree. This can be deduced by equating the transition probability in Equation 4 with the probability that the next coalescent will be in the minority clade, which is the ratio of the coalescent rate in the minority tree over the sum of coalescent rates in both the minority and majority trees.

Simulation of 100 genealogies from a discrete-event birth-death process has been previously described (McCloskey and Poon 2017; Vaughan and Drummond 2013). These simulations were based on a process with heterogeneous classes of individuals with different birth rates. With some probability, lineages migrate to a class with higher birth rates. This could represent a generic outbreak scenario such as a set of individuals with higher risk behaviour or other exposures. In a separate set of simulations, the outbreak population differs from the main population along multiple dimensions: the birth rate and the sampling rate are both increased by a common factor (5×). 100 genealogies were simulated under both scenarios and the *treestructure* algorithm was applied to each. To create more challenging conditions for the method and to evaluate the sensitivity of the method to sample coverage, we also applied the method to genealogies based on subsampled lineages with a frequency of 25%. Complete descriptions of parameters and simulation methods can be found in (McCloskey and Poon 2017).

The performance of *treestructure* was evaluated using the normalised mutual information (NMI) statistic and adjusted Rand index (ARI) computed using the *aricode* R package (Vinh et al. 2010). Both statistics quantify the strength of association between the estimated and actual structure of the tree, with larger values corresponding to higher quality reconstructions.

## Results

### Simulation studies

The *treestructure* algorithm achieves relatively high fidelity of classifications in comparison to other methods in the structured coalescent simulations which included 20% of samples from a rapidly growing outbreak. Figure 3 compares the values of NMI and ARI for three methods of structure analysis. In these statistics, the partition of the tree computed by each method is compared to the true membership of each sampled lineage in outbreak or in the constant-size reservoir population. Across 100 simulations, treestructure has mean ARI of 41% (IQR: 20-57%). The FastBAPS method (Tonkin-Hill et al. 2019) has mean ARI of 2.3% (IQR:1.2-3.3%) and the CLMP method (McCloskey and Poon 2017) has mean ARI 5.2% (IQR:-1-7.5%). The NMI statistic gives similar differences between the methods to ARI (Fig. 3).

**Figure 3:**
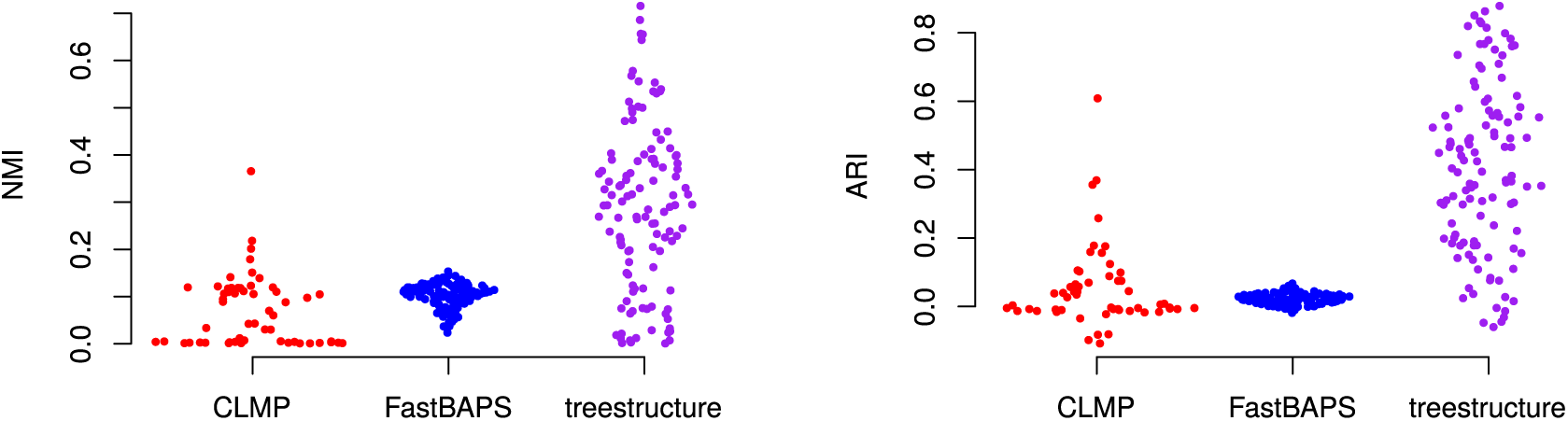
The normalised mutual information (NMI) and adjusted Rand index (ARI) as a function of classifications from several tree-partitioning algorithms and membership of lineages in outbreaks or a constant-size reservoir. Each point corresponds to a structured coalescent simulation where 20% of tips are sampled from an exponentially growing outbreak.

The lower performance of CLMP and FastBAPS in these comparisons is largely a consequence of false-positive partitioning of samples from the reservoir population, but CLMP and FastBAPS usually correctly identify a clade that closely corresponds to the outbreak. In contrast, the *treestructure* method seldom sub-divides clades from the reservoir. Figure 4 compares the entropy of partition assignments only within lineages sampled from the outbreak. This shows that all methods are assigning outbreak lineages to a small number of partitions and no method is clearly superior by this metric. The CLMP method has the lowest entropy (mean 0.40) but also several large outliers. *treestructure* has higher entropy (mean 0.57) but few outliers. FastBAPS has even higher entropy (mean 0.68) with a long tail of high values (Fig. 4).

**Figure 4:**
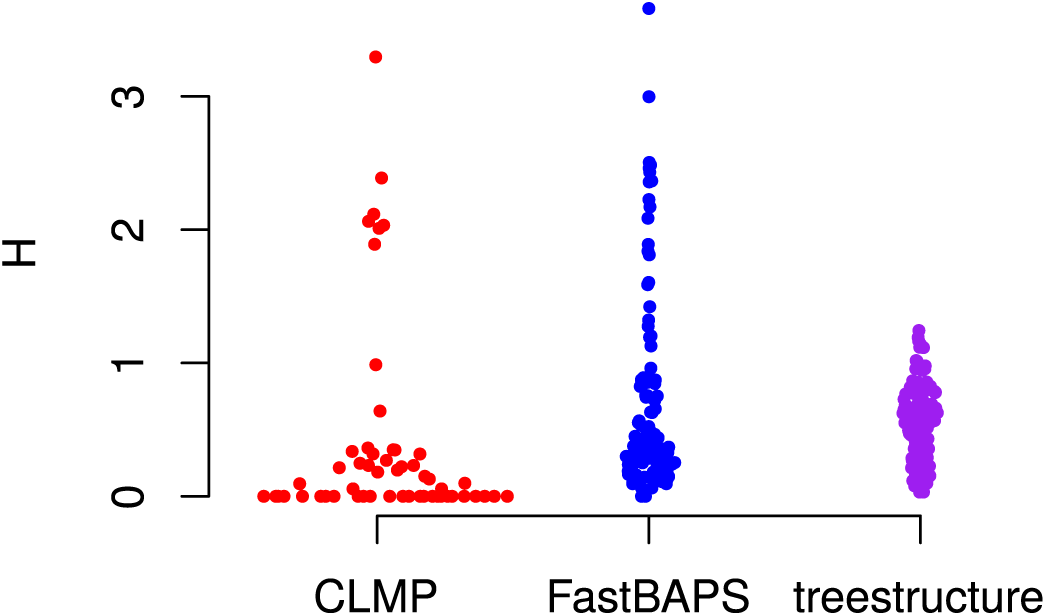
Entropy (*H*) of classification from several tree partitioning algorithms applied to the structured coalescent simulations but only counting lineages sampled from the exponentially growing outbreak.

The performance of all methods depended on the sample density and growth rate of the outbreak. Fast growing outbreaks are easier to detect by all methods but the role of sample density is more ambiguous. The Pearson correlation of ARI with growth rate is 53%, 71% and 27%, for *treestructure*, FastBAPS, and CLMP respectively. Not all methods are equally sensitive to these parameters however and FastBAPS is especially sensitive to growth and sample density. The growth rate and sample density collectively explain 41%, 60%, and 28% of variance of ARI in *treestructure*, FastBAPS, and CLMP respectively.

We also performed analyses with Phydelity, a recently proposed method for transmission cluster identification (Han et al. 2018). This tended to generate a very large number of clusters, both within and outside of the outbreak demes, reflecting a different emphasis of this method on finding closely related clusters rather than addressing differences in macro-level population structure. Thus, results with Phydelity and other clustering methods were not easily comparable to *treestructure*.

Figure 5 shows performance of *treestructure* on previously published tree simulations (McCloskey and Poon 2017). These simulations differ from the structured coalescent simulations presented above because both the reservoir and outbreak demes are growing exponentially at different rates. The birth rate in the outbreak deme is five-fold the birth rate in the reservoir, but in one set of simulations, both the birth rate and sampling rate in the outbreak was also increased five-fold. In these simulations, the performance of *treestructure* (mean ARI 53%) is slightly lower than the CLMP method (McCloskey and Poon 2017) (mean ARI 72%) when only the birth rate differs in the outbreak deme. However *treestructure* maintains good performance when death and sampling rates also differ. In that case, *treestructure* has mean ARI 42% and CLMP has mean ARI 0%. The results are similar when using NMI instead of ARI (Supplementary Fig. S1). The difficulty of detecting outbreaks with different sampling patterns was previously highlighted as a challenge for CLMP (McCloskey and Poon 2017).

**Figure 5:**
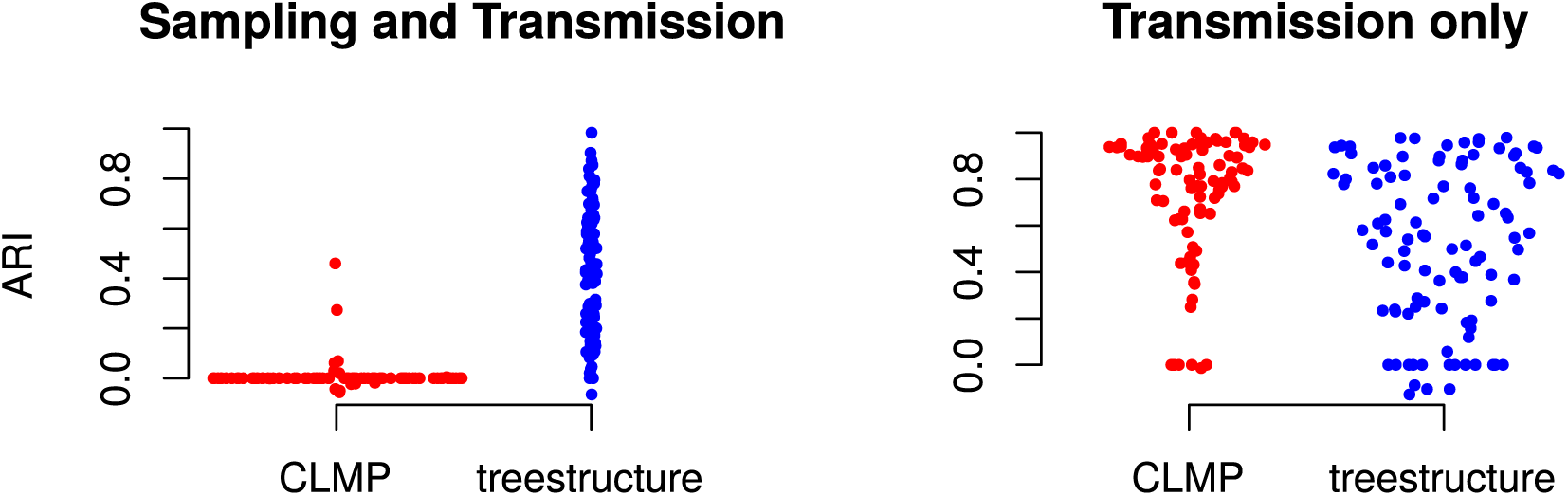
The adjusted Rand index for 100 previously published simulations (McCloskey and Poon 2017). This describes accuracy of classification of tips into outbreaks using the *treestructure* method and CLMP. Results on the left were based on simulations where both transmission and sampling rates varied in the outbreak cluster, whereas simulations on the right only allowed transmission rates to vary.

Simulations of unstructured Kingman coalescent trees shows that the distribution of the standardized test statistic is approximately normal (Supplementary Fig. S2). The quality of the normal approximation depends on the extent of phylogenetic error. In estimated phylogenies based on simulated sequence data, there is substantial skew in the test statistic which is most pronounced for larger clades that have a more distant MRCA (Supplementary Fig. S3). The extent of error due to phylogeny estimation will depend on many variables as well as on the choice of methodology when estimating time-scaled trees; in this case, effective population size and substitution rates were chosen to yield a data set with comparable diversity to a real HIV sequence data set, and there is considerable error in the estimated date of the TMRCA and tree topology which was estimated using the neighbor joining method. In the absence of phylogenetic error, the false positive rate based on a 95% confidence threshold was 5.1%. With phylogenetic error, the false positive rate increased to 12.2%.

Analysis of trees simulated with predefined structure showed that statistical power increases as expected with sampling density and effective population size contrast between the two clades. Supplementary Figure S4 shows the normalized test statistic for various sample sizes and contrasts of effective population size in two clades descended from the root of a tree. The statistic significantly deviates from zero with increasing sample sizes and with increasing differences in effective population sizes. For example, using a 95% confidence level, we find a significant difference between clades in 85% of simulations sampling 40 tips from the minority clade and with a two-fold difference in the rescaled effective population sizes. This decreases to 40% of simulations if sampling only 10 tips, but increases to 100% if there is a five-fold difference in the scaled effective population sizes.

### Clonal expansion of drug-resistant *N. gonorrhoeae*

We examined the role of evolution of antimicrobial resistance in shaping the phylogenetic structure of *N. gonorrhoeae* using 1102 previously described whole genome sequences (Grad et al. 2016). These isolates were collected from multiple sites in the United States between 2000 and 2013 and featured clonal expansion of lineages resistant to different classes of antibiotics. We estimated a maximum likelihood tree using *PhyML* (Guindon et al. 2010) and corrected for the distorting effect of recombination using *ClonalFrameML* (Didelot and Wilson 2015). We estimated a rooted time-scaled phylogeny using *treedater* (Volz and Frost 2017). A relaxed clock model was inferred, with a mean rate of 4.6 × 10^−6^ substitutions per site per year. *BactDating* (Didelot et al. 2018) was also applied for the same purpose and found to give very similar estimates for the clock rate and dating of clades.

We focus on the origin and expansion of two clades which independently developed resistance to cefixime (CFX) by acquiring the mosaic *penA* XXXIV allele (Grad et al. 2016). Note, however, that the level of susceptibility to CFX varies, particularly in the largest of these two clades. In one lineage within this clade, the mosaic *penA* XXXIV allele was replaced by recombination with an allele associated with susceptibility. Other isolates within this clade gained mutations that further modified the extent of resistance. The largest of the two clades emerged on a genomic background that was already resistant to ciprofloxacin (CIP), so that it has reduced susceptibility to both CIP and CFX. The smallest of the two clades is resistant to CFX but not CIP. To further analyse the relationship between CFX resistance and N. gonorrhoeae population structure, we focused our analysis on a tree with just 576 tips, representing the genomes from these two CFX resistant clades as well as genomes from the two clades that are most closely related to the two CFX resistant clades. The output of *treestructure* is shown in Figure 6, using unique colours to highlight each of the 11 clusters that were identified with *α* = 1%. The clusters reported by *treestructure* are highly correlated with CFX resistance. Among all distinct pairs of sampled isolates, 84% share the same resistance profile and cluster membership.

**Figure 6:**
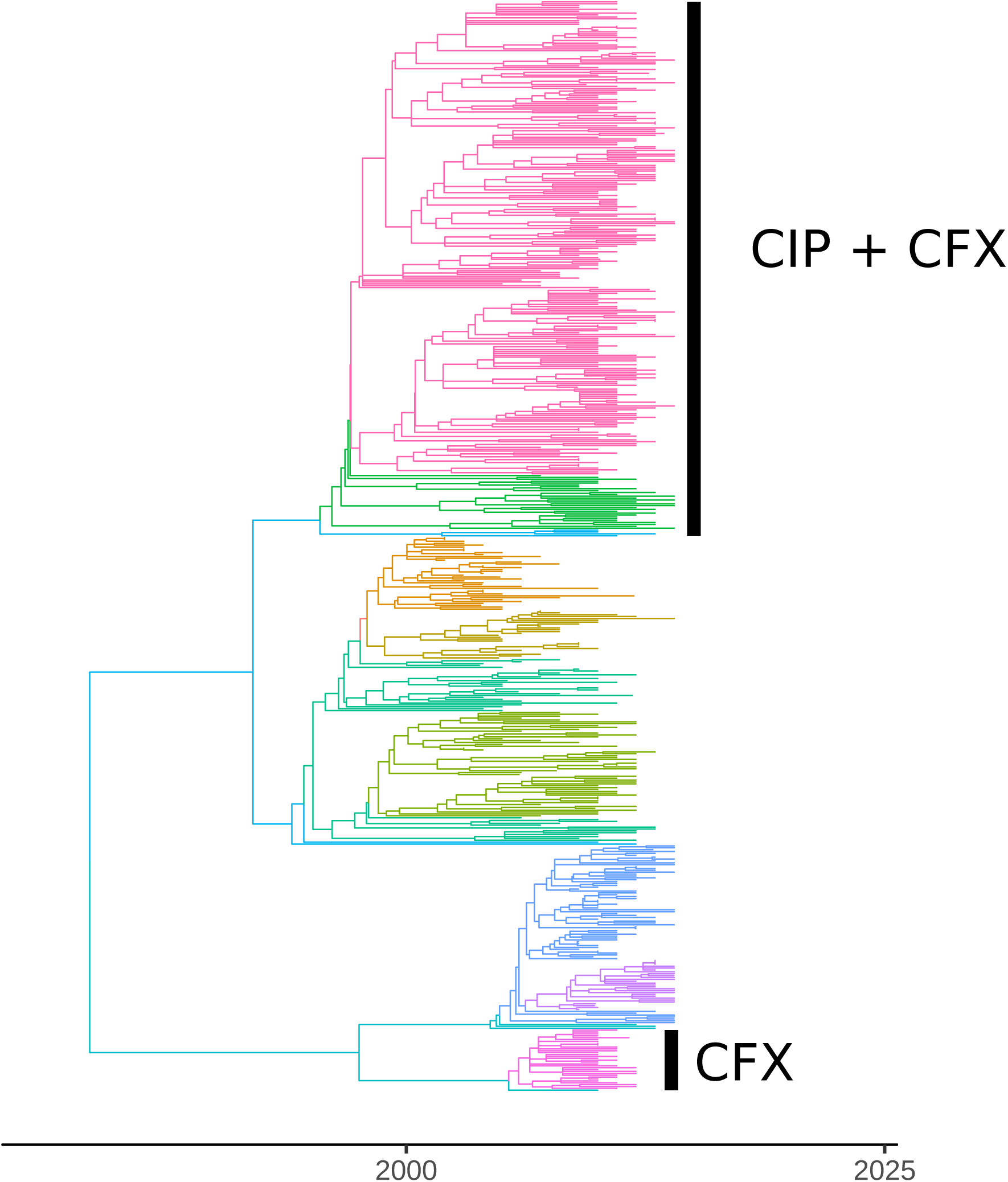
A time-scaled phylogeny based on 576 whole genomes of *N. gonorrhoeae*, comprising two clades with reduced susceptibility to cefixime (CFX) and their two sister clades. The top clade also has resistance to ciprofloxacin (CIP). Different colours on the tree represent the partition detected using the *treestructure* algorithm.

We compared *treestructure* with a different method for detecting community structure, FastBAPS (Tonkin-Hill et al. 2019), since BAPS models are often applied to bacterial pathogens. We applied FastBAPS using the same time-scaled phylogeny described previously and using a trimmed sequence alignment consisting of 38830 polymorphic sites and removing sites with many gaps. This produced a similar partition of the tree (Supplementary Fig. S5) with a few differences. The FastBAPS clusters overlap exactly with the clade featuring dual resistance (CIP and CFX), whereas *treestructure* classified a small number of deep-splitting lineages into a different cluster. Note however that this behaviour is not necessarily problematic, and may represent a progressive increase in fitness following the acquisition of resistance through the evolution of compensatory mutations (Didelot et al. 2016). Indeed, we found a significant difference in the resistance profile of the two *treestructure* clusters within the clade resistant to both CIP and CFX: the smallest cluster had a greater frequency of high resistance to CIP compared to the largest cluster (100% and 81%, respectively).

FastBAPS did not identify the smaller clade with resistance to CFX and not CIP and instead grouped that clade with its sensitive sister clade. In general, *treestructure* found many more clusters within the two sister clades and FastBAPS tended to group these together. We also applied the much more computationally intensive RhierBAPS method (Tonkin-Hill et al. 2018), and obtained almost identical results to FastBAPS. Overall, BAPS methods appear to give more weight than *treestructure* to long internal branches when identifying clusters.

### Epidemiological transmission patterns of HIV-1

We reanalysed a time-scaled phylogeny reconstructed from 2068 partial *pol* HIV-1 subtype B sequences collected from Tennessee between 2001 and 2015 (Dennis et al. 2018). Each lineage within this phylogeny corresponds to a single HIV patient sampled at a single time point, and various clinical and demographic covariate data concerning these patients can be associated with each lineage. In the original study, these sequence data were used to show high rates of transmission among young (age *<* 26.4 years old) men who have sex with men (MSM) (Dennis et al. 2018). Clustering by threshold genetic distance is often used in HIV epidemiology (Dennis et al. 2014) and indicated that young white MSM had the highest odds of clustering.

We applied the *treestructure* algorithm with default settings to the time-scaled tree which yielded ten partitions with sizes ranging from 58 to 398. The tree and partitions are shown in Figure 7 where partitions are labeled according to the median year of birth among patients in each partition. Many of these partitions were polyphyletic, suggesting possible multiple importations of lineages to specific risk groups. We then compared the estimated partition of the tree with patient covariates. A particular partition stands out along multiple dimensions: it is the smallest (size 58), polyphyletic, arose in the recent past, and is characterised by very young MSM. The median year of birth in this partition is 1987, in stark contrast to the rest of the sample with year of birth in the 1970s. Clades within this young partition are also nested paraphyletically under other relatively young partitions (Fig. 7).

**Figure 7:**
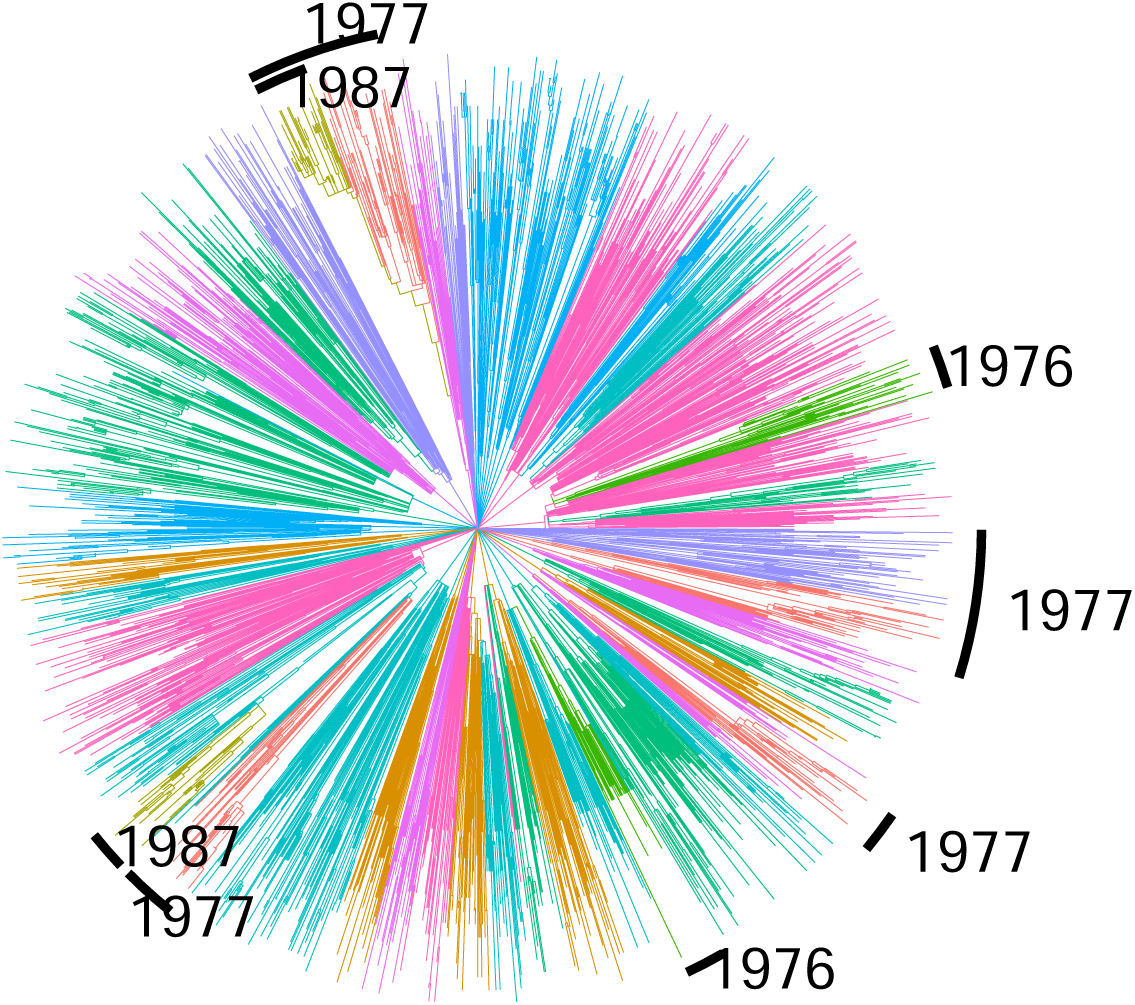
A time-scaled phylogeny estimated from HIV-1 *pol* sequences in Tennessee (Dennis et al. 2018). The colours correspond to the ten partitions identified using the *treestructure* algorithm. Several partitions are annotated with the median year of birth of HIV patients from whom sequences were sampled. Unannotated partitions had years of birth 1969-1972.

We did not find a significant association between the tree partition and residential postal codes (Tukey analysis of variance, *p* = 0.097). This is in agreement with the original study which found minimal impact of geography on genetic clustering in this sample, however this is largely a consequence of the highly concentrated nature of the sample around Nashville. The ethnicity of patients (black, white, and other) was strongly associated with the estimated partition. Black MSM were strongly concentrated in the 1987 partition in particular (83% in contrast to 26-38% in all other partitions). The odds ratio of black ethnicity given membership in the 1987 partition was 9.7 (95% CI:5.2-19.8).

Finally, we performed a phylodynamic analysis to investigate if the partition structure supported the previously published findings that young MSM were transmitting at a higher rate (Dennis et al. 2018). To estimate the temporal variations in the effective population size, we used the nonparametric *skygrowth* R package (Volz and Didelot 2018). We estimated *N*_*e*_(*t*) for each partition individually using a range of precision parameters which control the smoothness (*τ*) of the estimated trajectories since we lack a priori information about volatility of these trajectories. Figure 8 shows *N*_*e*_(*t*) for each partition with *τ* = 10 and Supplementary Figures S6 and S7 show results using different values of *τ*. The 1987 partition again stands out as the only group which shows evidence of recent and rapid population growth. Less dramatic recent periods of growth are also noticeable for other partitions with young patients. The current exponential growth in the 1987 partition is not consistent across all analyses, but when *τ <* 10 we find *N*_*e*_(*t*) drops precipitously in 2014-2015 (Supplementary Fig. S6). However, this could also be an artefact of non-random sampling and inclusion of transmission pairs within the sample.

**Figure 8:**
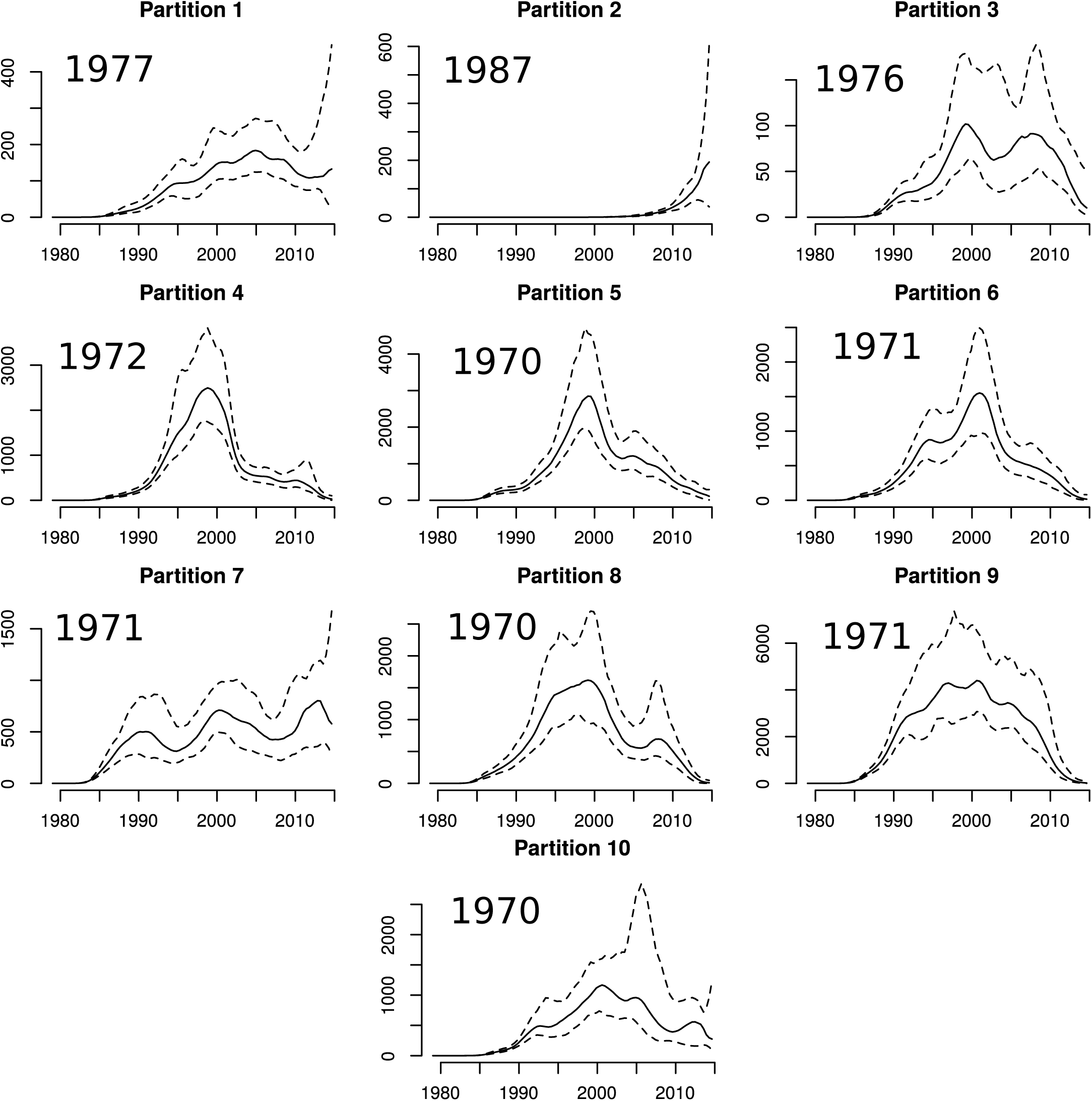
Estimated effective population size through time for each partition in the Tennessee HIV-1 phylogeny. Each panel is annotated with the median year of birth among HIV patients in each partition. *N*_*e*_(*t*) was estimated using the *skygrowth* method (Volz and Didelot 2018) with precision parameter *τ* = 10.

This analysis supports the hypothesis that there has been a recent and rapid increase in HIV transmissions among young MSM in Tennessee and in particular among young black MSM. This interpretation is mostly in agreement with the original study (Dennis et al. 2018), but we find that black MSM are a group at greater risk than young white MSM.

## Discussion

Contrasting the distribution of ordering of nodes provides a natural criterion for distinguishing clades within a time-scaled phylogeny which are shaped by different evolutionary or demographic processes. The non-parametric nature of this classification method imposes minimal assumptions on the mechanisms that generate phylogenetic patterns. Thus, we have found this method maintains good performance over a diverse range of situations where phylogenetic structure is produced, including differential transmission rates, epidemiological outbreaks, evolution of beneficial mutations, and differential sampling patterns. Our work is related to the research on species delimitation methods (see for example Zhang et al. 2013) although targeted at within-species variation, and is also related to recent work on methods for detecting co-diversification of species (Oaks et al. 2019). This method appears relatively robust compared to other methods against false-positive identification of phylogenetic structure, but nevertheless has good sensitivity for detecting structure in most situations.

There are many immediate applications of this method in the area of pathogen evolution where time-scaled phylogenetics is increasingly used in epidemiological investigations (Biek et al. 2015). We have demonstrated the role of selection in shaping phylogenetic structure of *N. gonorrhoeae*, and our method clearly identifies clades which expanded in the recent past due to acquisition of antimicrobial resistance. We have demonstrated the role of human demography and transmission patterns in shaping the evolution of HIV-1, and our method has shown distinct outbreaks of HIV-1 in specific groups defined by age, race, and behaviour. Furthermore, we have shown how clades detected by this method can be analysed using phylodynamic methods that can yield additional insights into recent outbreaks or the mechanisms which generated phylogenetic structure. For example, we have applied non-parametric methods to estimate the effective population size through time in HIV outbreaks detected using *treestructure* which highlighted particular groups that appear to be at higher risk of transmission. Such analyses would be more problematic using other partitioning or clustering algorithms because phylogenetic clusters can appear by chance in homogeneous populations of neutrally evolving pathogens, and this can give the false appearance of recent growth (Dearlove et al. 2017). This application of phylodynamics analysis methods is possible because the statistical test used in *treestructure* provides theoretical justification for treating each partition as a separate unstructured population.

Applications of the *treestructure* algorithms are scalable to relatively large phylogenies. The main algorithms require only a single pre-order traversal of the tree and all of the computations presented here required less than one minute to run. The method is based on a time-scaled phylogeny, and the computational burden of this preliminary step is typically higher than that of running *treestructure*, even though significant progress has been made recently in this area (Volz and Frost 2017; Didelot et al. 2018; Sagulenko et al. 2018; Tamura et al. 2018; Miura et al. 2019). Future developments of *treestructure* and other methods post-processing time-scaled phylogenies (Volz and Didelot 2018; Didelot et al. 2017) should address the uncertainty in the input phylogeny, for example by accounting for bootstrap or Bayesian support values for phylogenetic splits, or by summarising results from multiple trees.

## Algorithm 1: Algorithm for computing the null distribution and associated p-value of the test-statistic for cladistic outliers.

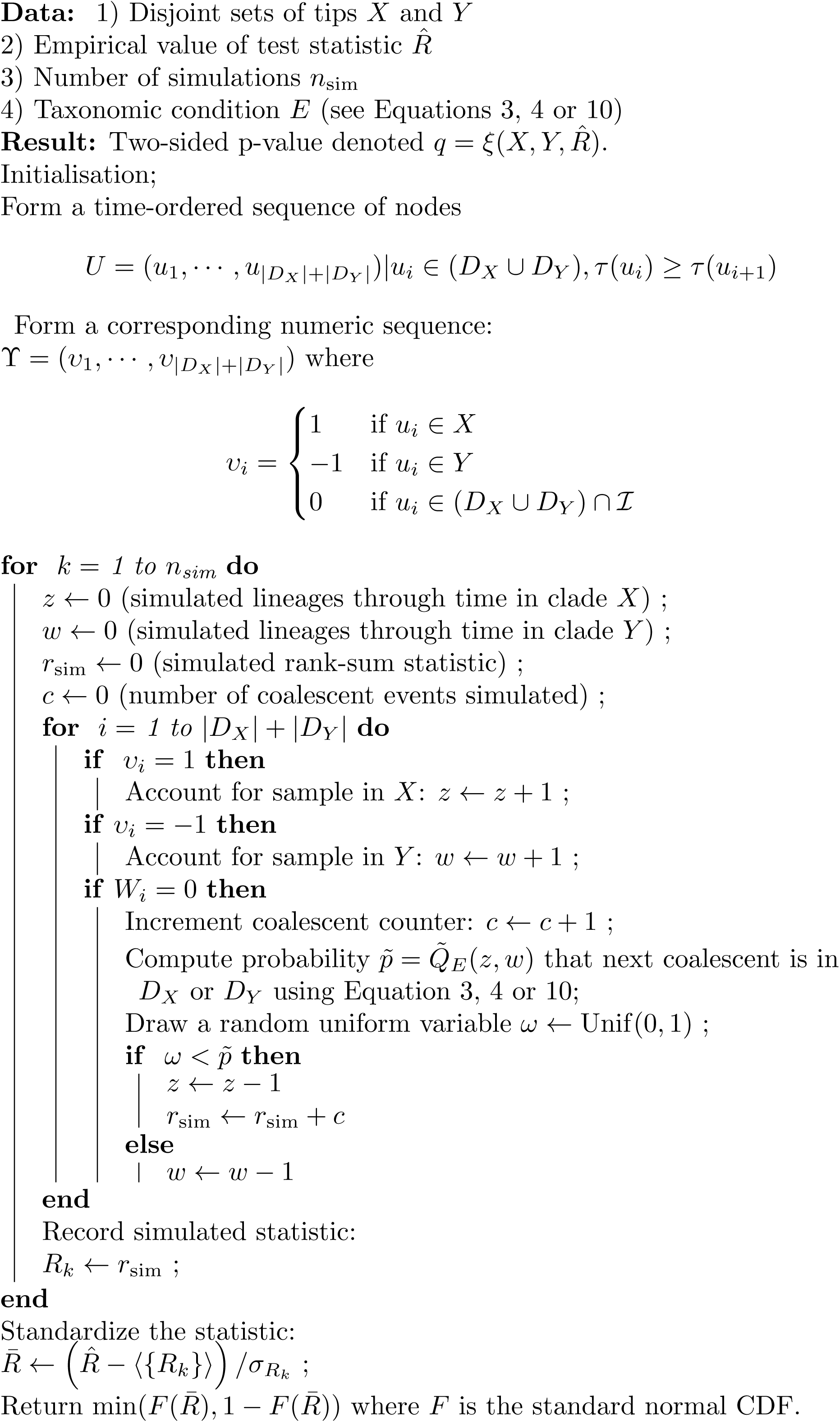

## Algorithm 2: Algorithm for detecting cladistic outliers.

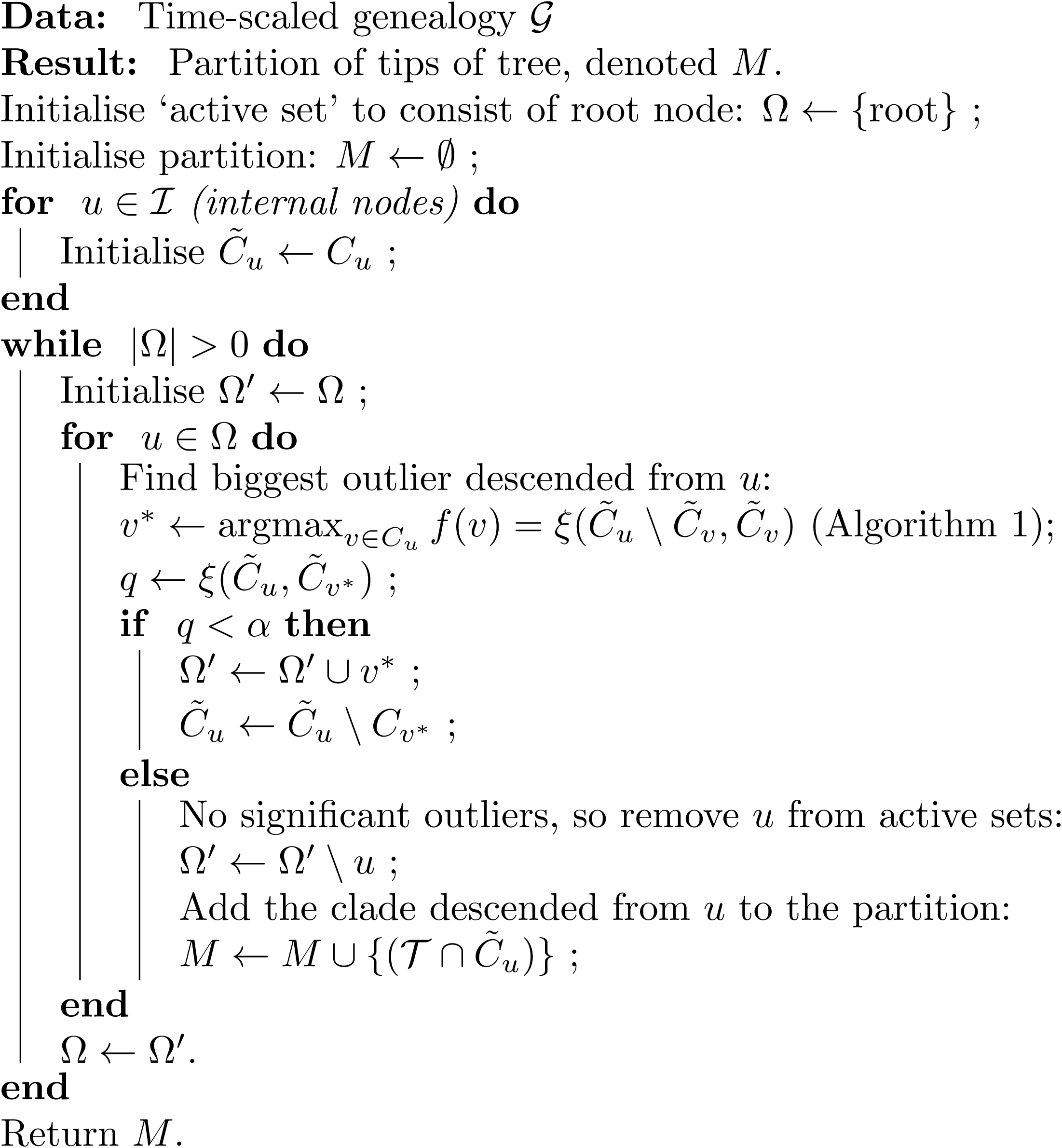

**Figure S1:**
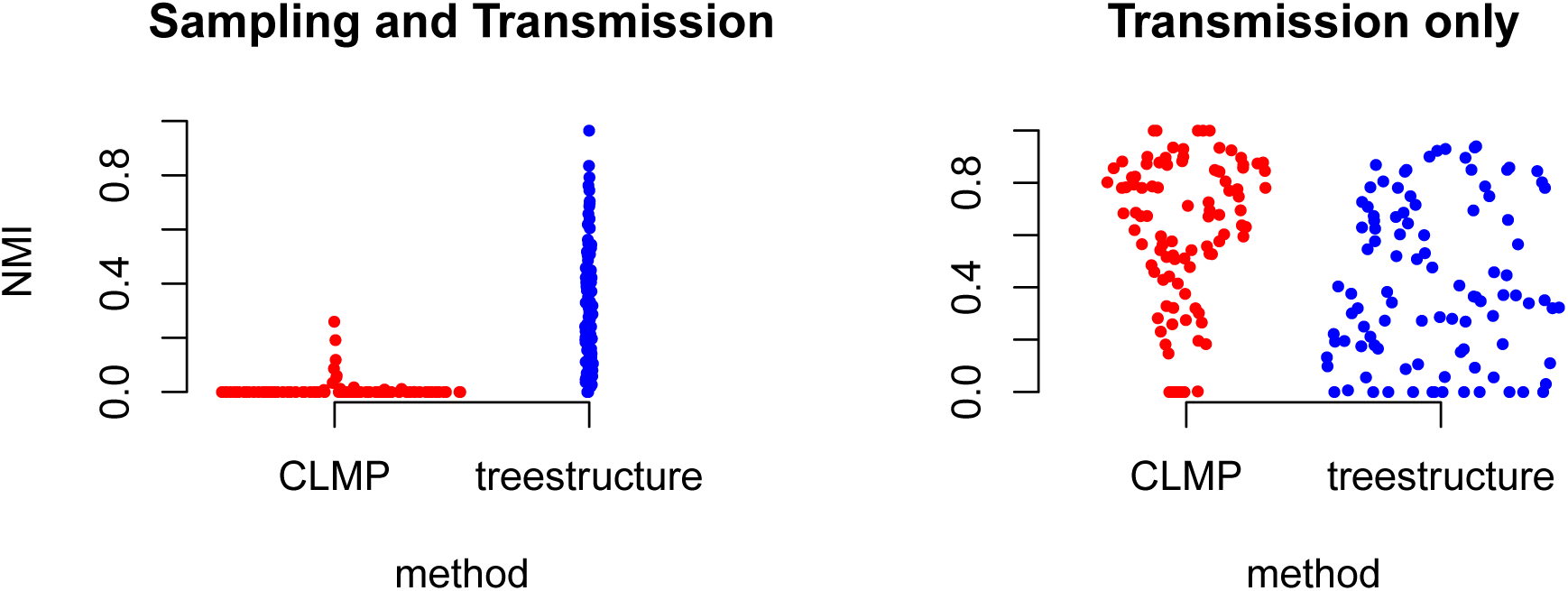
The normalised mutual information (NMI) for 100 previously published simulations (McCloskey and Poon 2017). This describes accuracy of classification of tips into outbreaks using the *treestructure* method and CLMP (McCloskey and Poon 2017). Results on left were based on simulations where both transmission and sampling rates varied in the outbreak cluster, whereas simulations on the right only allowed transmission rates to vary.

**Figure S2:**
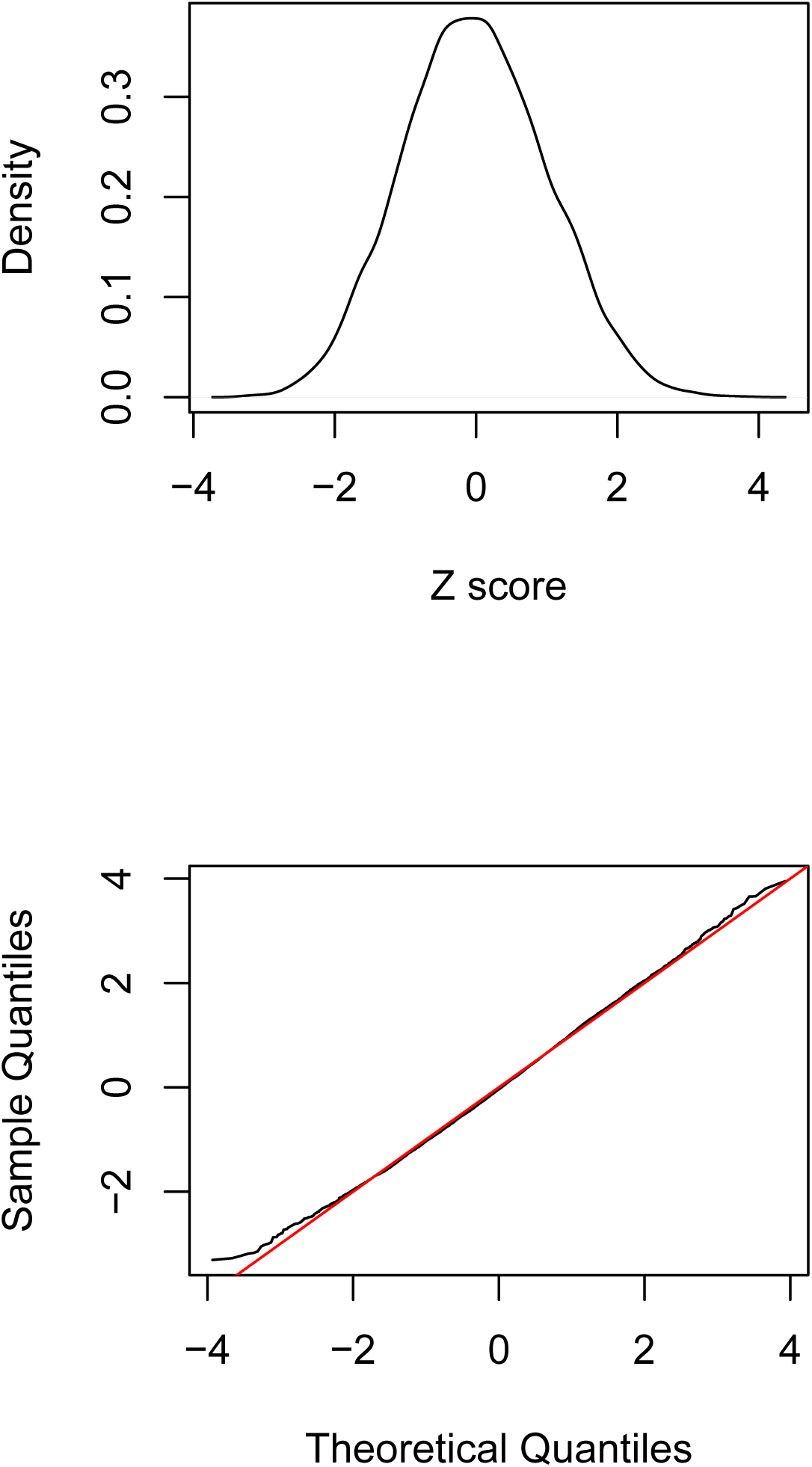
The distribution of the test statistic under the null hypothesis with Kingman coalescent trees simulated with 50 tips. Top: The empirical density of the standardized test statistic (Z score) across internal nodes in 1,000 Kingman coalescent trees. Bottom: A quantile-quantile plot of the Z scores from internal nodes in 1,000 coalescent trees and the standard normal distribution.

**Figure S3:**
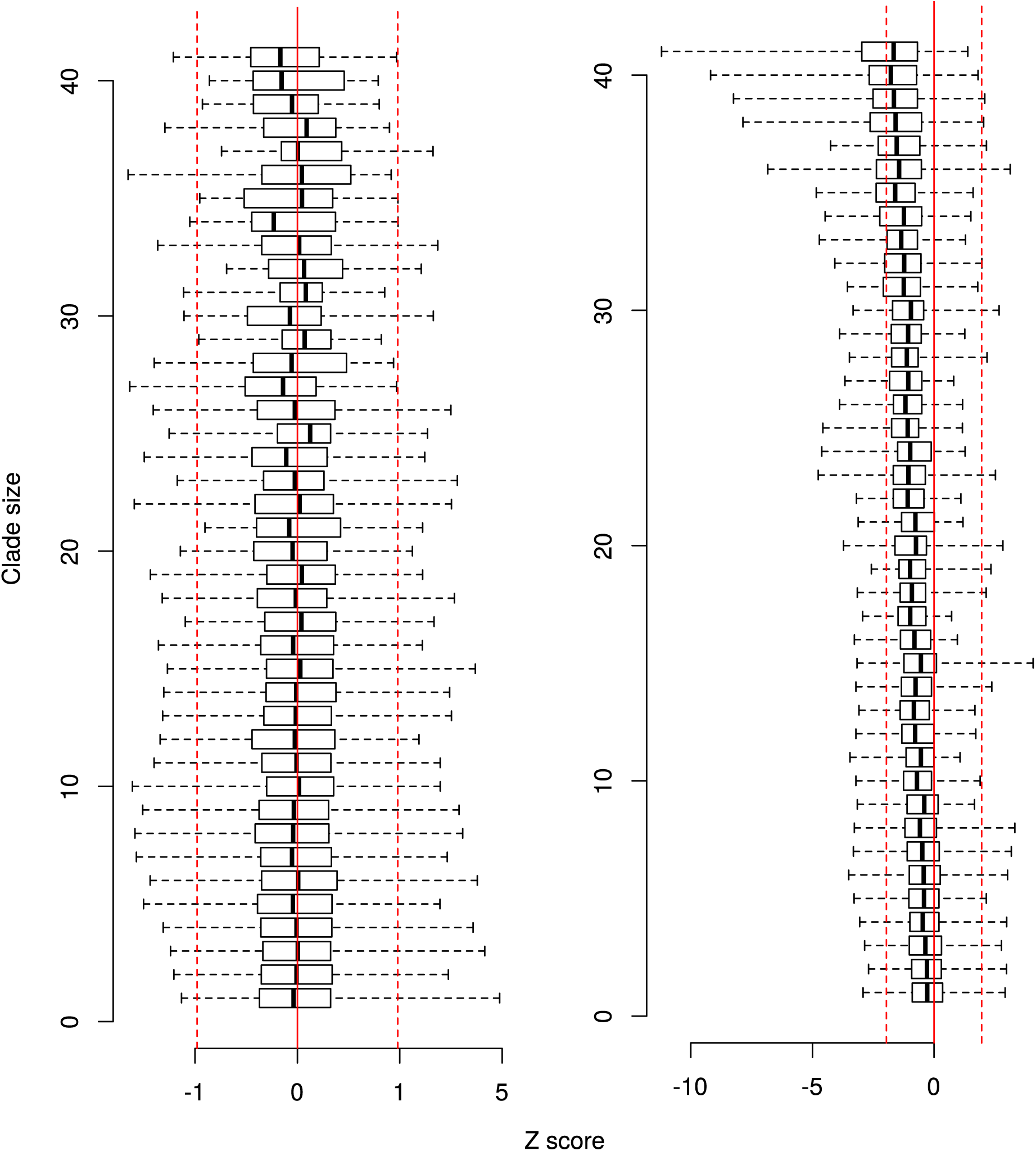
Distribution of the standardized test statistic (Z scores) under the null hypothesis and tabulated by clade size. Each box shows the range (whisker) and interquartile range (box) of Z scores across 1,000 simulated coalescent trees and for a particular clade size (number of tips). The red lines show the interval corresponding to a 95% confidence region. The left part is based on Kingman coalescent trees, while the right part is based on estimated time-scaled phylogenies using simulated sequences as described in the text.

**Figure S4:**
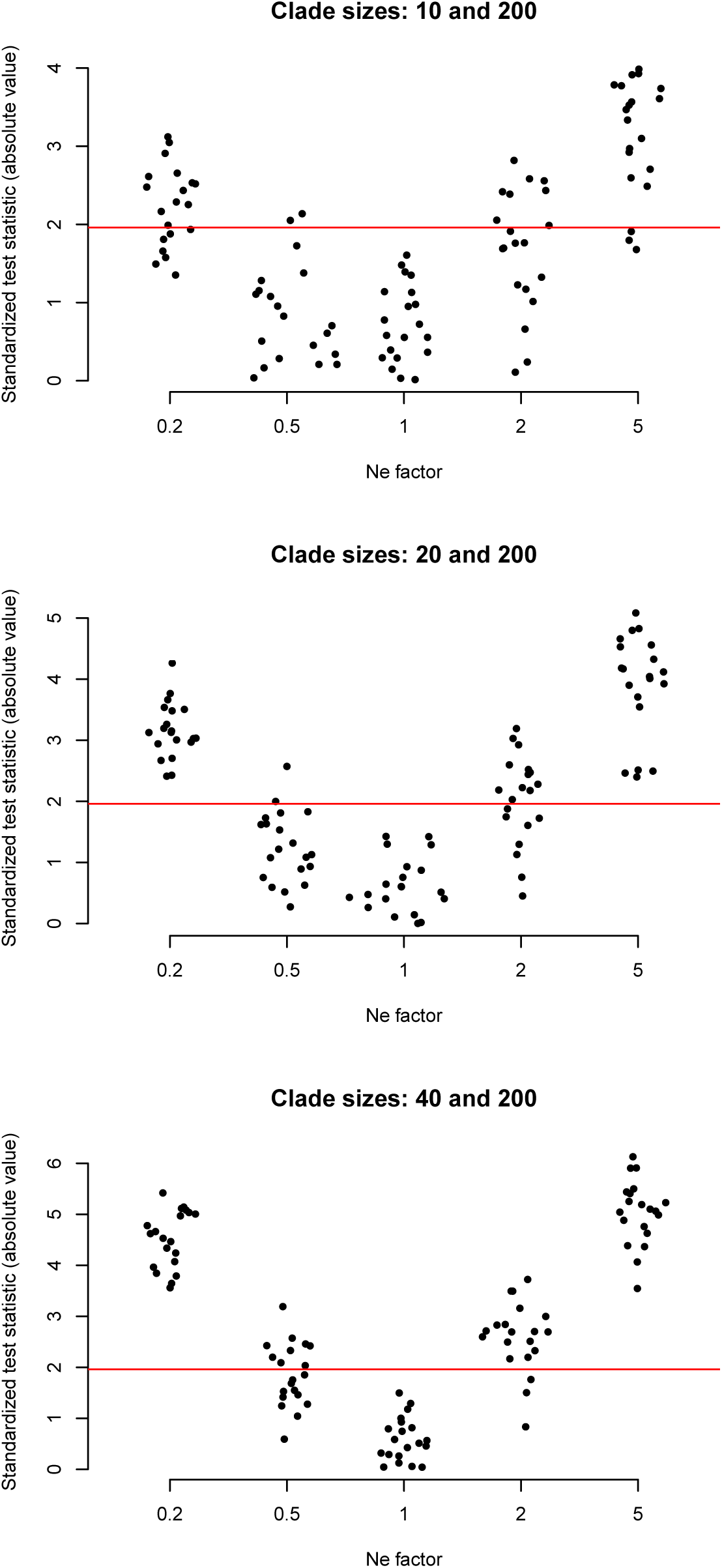
Power to discriminate between clades as a function of sample size and difference in effective population size. Each plot shows the absolute value of the standardized test statistic of the MRCA of a minority clade. The minority clade has an effective population size selected to provide various levels of contrast with the majority clade (see text). The x-axis shows 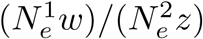 where *z* and *w* are the number of tips in the minority and majority clades, and 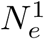 and 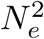 are the effective population sizes in the minority and majority clades. The red line corresponds to 1.96 which is the 95% quantile of the standard normal distribution. The top, middle and bottom panels are each based on simulations where the minority clade had 10, 20, and 40 tips respectively, whereas the majority clade always had 200 tips.

**Figure S5:**
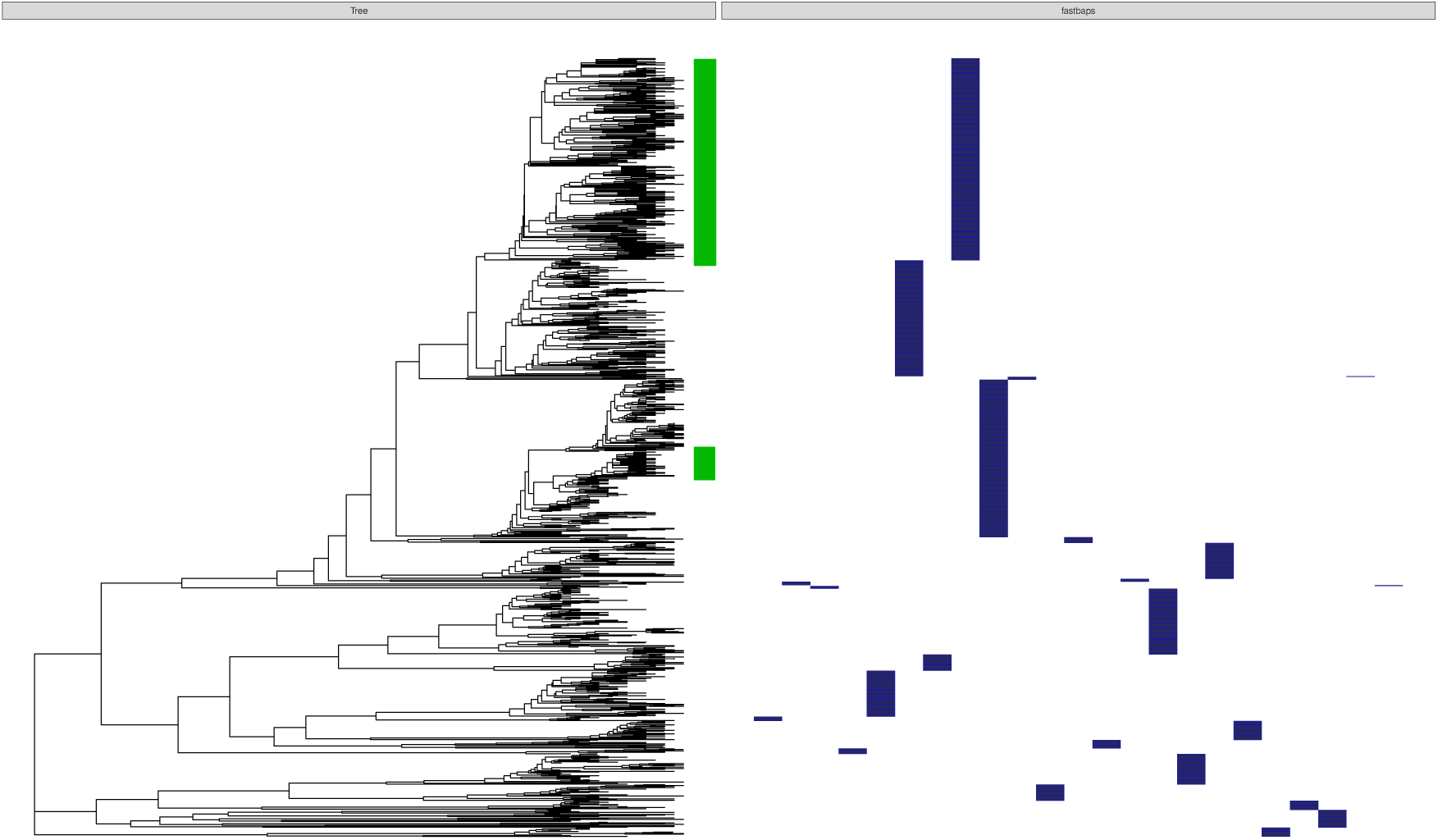
The output of FastBAPS classification applied to 1102 *N. gonorrhoeae* isolates described in the main text. Clades indicated in green have CFX resistance.

**Figure S6:**
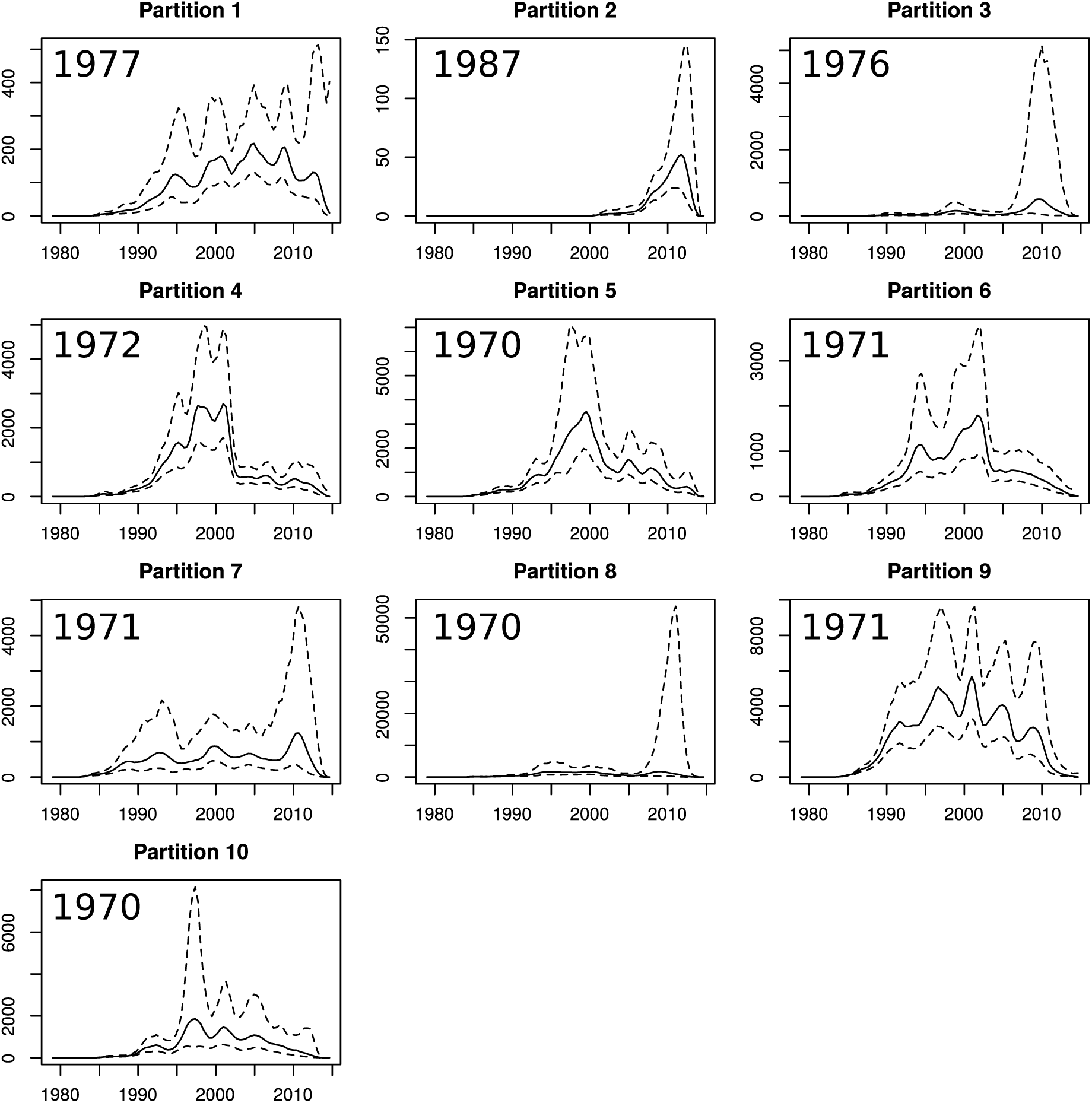
Estimated effective population size through time for each partition in the Tennessee HIV-1 phylogeny. *N*_*e*_(*t*) was estimated using the *skygrowth* method (Volz and Didelot 2018) with precision parameter *τ* = 1.

**Figure S7:**
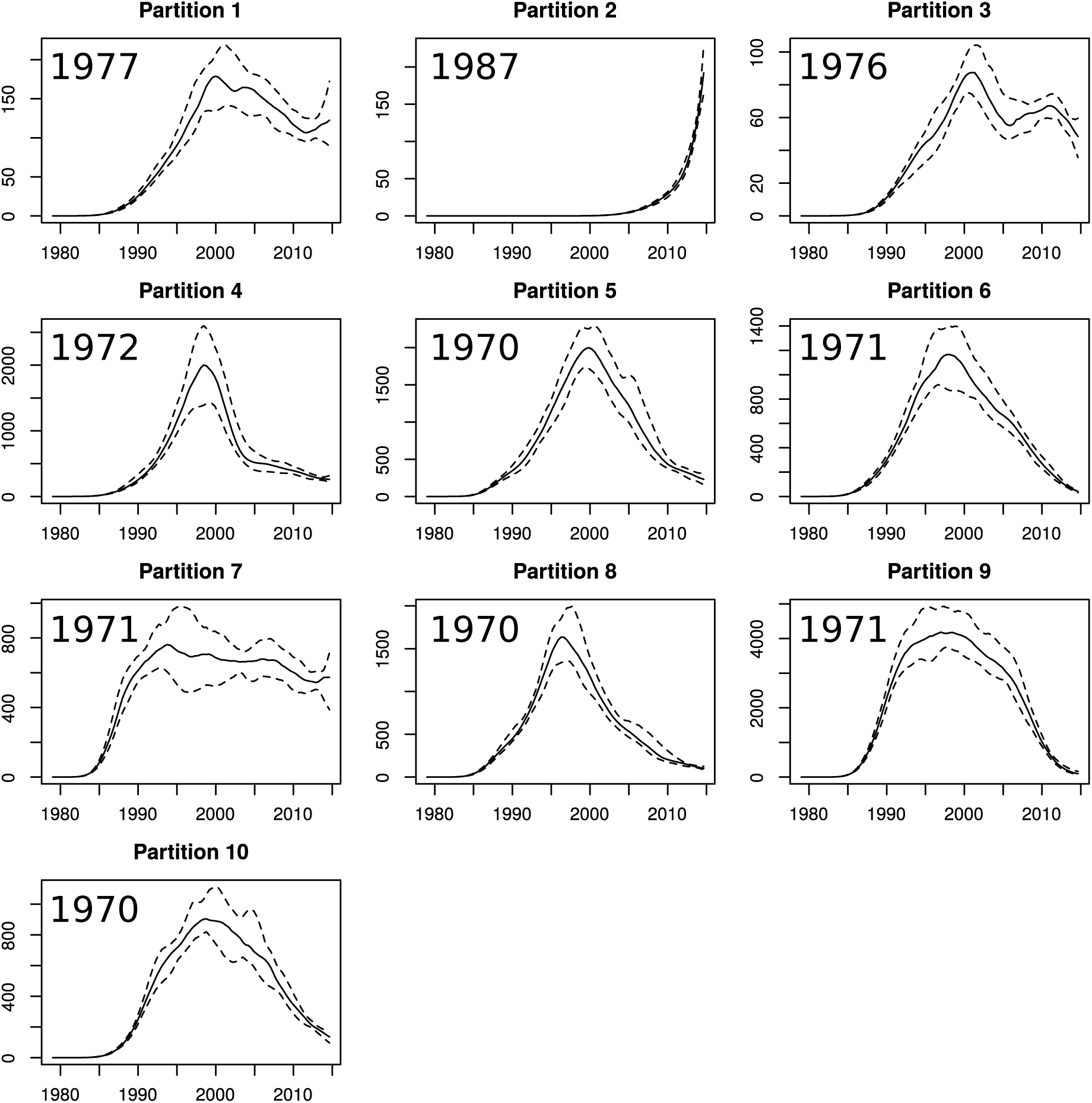
Estimated effective population size through time for each partition in the Tennessee HIV-1 phylogeny. *N*_*e*_(*t*) was estimated using the *skygrowth* method (Volz and Didelot 2018) with precision parameter *τ* = 100.

